# Insecticide resistance status of malaria vectors *Anopheles gambiae s.l* of southwest Burkina Faso and residual efficacy of indoor residual spraying with microencapsulated pirimiphos-methyl insecticide

**DOI:** 10.1101/2020.07.13.200865

**Authors:** Dieudonné D. Soma, Barnabas M. Zogo, François D. Hien, Aristide S. Hien, Didier P.A. Kaboré, Mahamadi Kientega, Anicet G. Ouédraogo, Cédric Pennetier, Alphonsine A. Koffi, Nicolas Moiroux, Roch K. Dabiré

**Author notes:** These authors contributed equally to this work (Dieudonné D. Soma, Barnabas M. Zogo). These authors also contributed equally to this work (Nicolas Moiroux, Roch K. Dabiré).

## Abstract

The rapid spread of insecticide resistance in malaria vectors and the rebound in malaria cases observed recently in some endemic areas underscore the urgent need to evaluate and deploy new effective control interventions. A randomized control trial was conducted with the aim to investigate the benefit of deploying complementary strategies, including indoor residual spraying (IRS) with pirimiphos-methyl, in addition to long-lasting insecticidal nets (LLINs) in Diébougou, southwest Burkina Faso. We measured the susceptibility of *Anopheles gambiae s.l.* population from Diébougou to conventional insecticides. We further monitored the efficacy and residual activity of pirimiphos-methyl on both cement and mud walls using a laboratory susceptible strain (Kisumu) and the local *An. gambiae s.l.* population. *An. Gambiae s.l.* from Diébougou was resistant to pyrethroids (deltamethrin, permethrin and alphacypermethrin) and bendiocarb but showed susceptibility to organophosphates (pirimiphos-methyl and chlorpyrimiphos-methyl). A mixed-effect generalized linear model predicted that pirimiphos-methyl applied on cement or mud walls was effective for 210 days against the laboratory susceptible strain and 247 days against the local population. The residual efficacy of pirimiphos-methyl against the local population on walls made of mud was similar to that of cement (OR=0.792, [0.55-1.12], Tukey’s test p-value =0.19). This study showed that one round of IRS with pirimiphos-methyl CS has the potential to control the multi-resistant *An. gambiae s.l.* population from Southwest Burkina Faso for at least 7 months, regardless of the type of wall.

## Introduction

Current malaria vector control measures rely heavily on the use of Long Lasting Insecticidal mosquito Nets (LLINs) and indoor residual spraying (IRS) (World Health Organization [WHO] 2019a). Both strategies have had substantial impacts on malaria burden over the past 15 years. Indeed, LLIN and IRS accounted for an estimated 68 and 11% of the malaria averted cases, respectively, between 2000 and 2015 (Bhatt et al. 2015). Historically, IRS based on DDT was the cornerstone of the global malaria eradication campaign that has led to the elimination of malaria in 15 countries in Europe and America during the 1950s and 1960s. In Africa, however, these campaigns were not widely implemented due to a number of reasons including limited resources (WHO 1969). Subsequently, the coverage of IRS has dropped considerably in favor of LLINs. Until 2014, very few African countries still considered IRS as a prior action in malaria vector control (WHO 2015). More recently in 2017, IRS was implemented, either alone or in combination with LLINs in 40 African countries (WHO 2018a). Interest in combining IRS with LLIN seems to have increased in recent years across Africa because of the raise of pyrethroids resistance within the main major malaria vectors (WHO 2018b). As of 2017, the arsenal of insecticides recommended for IRS has been improved considerably making available 5 classes of insecticides including organochlorines, carbamates, organophosphates, pyrethroids and neonicotinoids (WHO 2011, 2019b). The Global Plan for Insecticide Resistance Management (GPIRM) recommends rotation of non-pyrethroid insecticides for IRS in areas where IRS and LLIN are combined (WHO 2012). However, options available for continued insecticide rotation are very limited in many endemic countries because resistance to multiple insecticide classes is very common in vector populations. According to WHO, resistance to organochlorines and carbamates were confirmed, respectively, in 62.4% and 30.6% of the sites tested in Africa between 2010 and 2016 (WHO 2018b). Resistance to organophosphate was less common, with 14.1% of the sites tested in Africa confirmed its occurrence (WHO 2018b).

In addition to the insecticide physiological resistance, a variety of factors can affect the effectiveness of IRS. Indeed, residual life and efficacy of the insecticide used can vary according to the formulation, the quality of sprays, and the type of walls (cements, mud, wood) (Sibanda et al. 2011, Djènontin et al. 2013).

We recently conducted a randomized-controlled trial in the rural area of Diébougou, Southwest Burkina Faso, to investigate whether the use of complementary strategies together with LLINs affords additional reduction in malaria transmission and cases. One of the strategies evaluated was IRS with microencapsulated formulation of pirimiphos-methyl. Microencapsulation is a technology that allows insecticides to last longer on substrates than usual (Poshadri and Kuna 2010). In the present study, we tested the susceptibility of the *An. gambiae s.l.* population from the rural area of Diébougou (southwest Burkina Faso) to conventional insecticides. Furthermore, we assessed the residual bio-efficacy of pirimiphos-methyl on mud and cement walls treated during the trial using a susceptible strain of *Anopheles gambiae s.s.* (*kisumu*) and a wild *An. gambiae s.l.* population.

## Materiels and Methods

### Study area

This study was carried out in two villages, Dangbara (−3.284°; 10.766°) and Nipodja (−3.383°; 10.988°), located in the Diébougou health district in southwest Burkina Faso. These villages were selected among the five villages which received a pirimiphos-methyl IRS intervention in a randomized control trial run in Diébougou, southwest Burkina Faso (Soma et al. 2019). The Diébougou area is characterized by an average annual rainfall of 1200 mm. The climate is tropical with two seasons: one dry season from October to May and one rainy season from June to September. Average daily temperature amplitudes are 18-36°C, 25-39°C and 23-33°C in dry cold (November to February), dry hot (March to May) and rainy season (June to October), respectively. Agriculture is the main economic activity in the area, followed by artisanal gold mining and coal and wood productions (Institut National de la Statistique et de la Démographie [INSD] 2015, 2017).

### Houses spraying

Actellic^®^300CS (Syngenta AG, Basel, Switzerland) was applied at a target dosage of one gram of active ingredient (pirimiphos-methyl) per square meter (1g a.i. /m^2^) in all houses of both villages in September 2017. We performed IRS using Hudson^®^ X-pert spray pumps (H.D. Hudson Manufacturing Company, Chicago, IL). The spray pumps (15 liters) were fitted with a 1.5 bar control flow valve on the lance pressure and equipped with ceramic 8002E nozzle according to WHO guidelines (WHO 2007). The spraying was performed by volunteers from the local communities who were trained by the National Malaria Control Program (NMCP) staff on a previous IRS campaign in Diébougou in 2012. We re-trained the spray operators and supervisors prior to IRS operations in the villages.

### Safety precautions

We took standard safety precautions with regard to mixing, handling and spraying insecticides (WHO 2006, 2007). Spray operators and supervisors used an appropriate protective equipment (gloves, hats, overalls, boots and facemasks). Spray operators, supervisors and householders were provided with an illustrated information sheet on the study, the possible adverse events in case of inappropriate spraying and safety precautions. We eliminated properly the leftover insecticides and bottles according to standard procedures (WHO 2007). The householders were advised by IRS operation teams about safety precautions in order to avoid possible risks during and after spray. They were advised to remain outside the rooms during spraying and until 3 hours after spraying. Adult heads of households were advised to ask their children not to intentionally touch the sprayed walls for at least one day after spraying, as the walls remained wet for about one day. We recommended the householders, if possible, to not scrub, mutilate or plaster the walls until the end of the study. The medical team of the Diébougou health district participated in the trial to attend to any medical illnesses of the inhabitants, or IRS operation team members.

### Chemical analysis

Before spraying, we attached Whatman No. 1 filter papers (10 x 10 cm) to the four inner walls of six randomly selected houses (three made of mud walls and three made of concrete walls) per village. On each four inner walls, two types of filters papers (one plasticized and one classical) were fixed in order to test for a possible migration of the insecticide from the filter papers to the wall as hypothesized by Moiroux *et al.* (2018). Plasticized and classical papers were fixed in areas where spray overlap was unlikely to ensure that the quantity of insecticide was constant. We also marked positions of filter papers on the wall to avoid carrying out subsequent cone bioassays at such surfaces. The filter papers were removed 24h after spraying and placed individually in aluminum foil with appropriate labels (village code, house number, type of surface and date of spraying). We stored the packed samples in a fridge at +4°C before sending them to the WHO collaborating center, Gembloux, Belgium for analysis of pirimiphos-methyl content.

### Insecticide resistance of wild *An. gambiae s.l* and residual efficacy

Both wild *An. gambiae s.l.* from the study area and the susceptible *An. gambiae s.s.* Kisumu strain (KISUMU1, MRA-762, VectorBase stable ID VBS0000026 on vectorbase.org) have been used in the following bioassay. We collected *An. gambiae s.l*. larvae in Bagane (3.150°; 10.575°, unsprayed village). Larvae were reared in the insectary of IRSS (temperature 27 ± 2 °C; relative humidity: 70±5%; 12h: 12h light: dark regimen) to adulthood. We fed larvae every day with Tetramin^®^ baby fish food. After emergence, mosquitoes were identified to species level using morphological keys (Gillies and Coetzee 1987). Adult *Anopheles* mosquito were provided with a sugar solution (10%), until their use for bioassays.

We tested the susceptibility of *An. gambiae s.l.* population from Bagane to 6 insecticides using the standard WHO protocol (WHO 2016). We exposed 4 replicate samples of 20-25 non blood-fed female, 3-5 days old, *An. gambiae s.l.*, for 60 min to each insecticides. We recorded mortality after 24 hours. Four insecticide classes were tested: carbamates (bendiocarb 0.1%), pyrethroids (alphacypermethrin 0.05%, permethrin 0.75% and deltamethrin 0.05%), organochlorine (DDT 4%) and organophosphates (chlorpyrifos-methyl 0.4% and pirimiphos-methyl 0.25%) (WHO 2016). As a negative control, two replicates of the same batch of mosquito were exposed to silicon oil impregnated papers. As a positive control, four replicates of susceptible *An. gambiae s.s.* Kisumu mosquitoes were tested with all insecticides.

In the same 12 houses randomly selected for chemical analysis, WHO cone tests were performed on days 2, 30, 60, 90, 120, 150, 180 and 210 post-spraying using both susceptible *An. gambiae s.s* and wild *An. gambiae s.l.* Bioassays were further performed on day 360, but only using the susceptible *An. gambiae s.s* Kisumu. In each house, we performed WHO cone tests on the four inner wall according to WHO guidelines (WHO 2016). A WHO cone test consists of introducing 10 to 15 unfed mosquitoes (3–5 days old) into a standard WHO cone for 30 minutes of exposure to the wall. As a control, four cone tests were performed on four unsprayed blocks. After exposure time, mosquitoes were placed in 150 mL plastic cups (1 replicate per cup) with 10% sucrose solution. All mosquitoes were held for 24h in the laboratory (27°C ±2°C and 70% ±5% relative humidity) to assess mortality.

### Statistical analysis

We analyzed insecticide susceptibility of the wild *An. gambiae s.l.* using a binomial generalized model with the mortality recorded in each tube as the response and the insecticide as fixed effect. The ‘brglm’ function of the ‘brglm’ package (Kosmidis 2019) in the software ‘R’(The R Development Core Team 2008) was used for this analysis. It allows to fit binomial-response regression models using the bias-reduction method developed by Firth (Firth 1993). These procedures return estimates with improved frequentist properties (bias, mean squared error) that are always finite even in cases where the maximum likelihood estimates are infinite (data separation). We used the ‘emmeans’ function of the ‘emmeans’ package to calculate estimated marginal means (EMM) of mortality for each insecticide and 95% confidence intervals (Russell et al. 2019).

We compared pirimiphos-methyl concentrations on filter papers using a linear (Gaussian) mixed effect models (LMM) with the wall surface (mud or cement), the type of filter paper (classical or plasticized) and interaction as fixed effect. The house and the wall in the house were set as nested random intercept. The Tukey’s post-hoc method was used to do multiple comparisons among modalities of the fixed terms (wall surface and paper type) using the ‘emmeans’ function of the ‘emmeans’ package (Russell et al. 2019). Mean differences (MD) and their 95% confidence interval were calculated.

For each strain, we analyzed mortality rate recorded in cones bioassays using a binomial response mixed effect model. We set the wall surface (cement or mud), the time after spraying (log-transformed) and interactions as fixed effects. The house was set as a random intercept. Odds-ratio (OR) and their 95% confidence intervals were computed. The ‘predict’ function in R applied on the bioassay mortality models was used to predict the time at which mortality fall under the 80% mortality threshold. We computed 95% confidence intervals of predictions.

### Ethics approval and consent to participate

The protocol of this study was reviewed and approved by the Institutional Ethics Committee of the Institut de Recherche en Sciences de la Santé (IEC-IRSS) and registered as N°A06/2016/CEIRES. Spay operators, supervisors and householders gave their written informed consent.

## Results

### Insecticide susceptibility bioassays

*An. gambiae s.l.* population from Diébougou were highly resistant to DDT and pyrethroids (alphacypermethrin, permethrin and deltamethrin), with mortality rates lower than 15%, as recorded in WHO susceptibility bio-assays (Fig. 1). This population was also resistant to bendiocarb (mortality rate = 67% Confidence Interval (CI) 95% [57; 75]). However, it was fully susceptible (100 % mortality) to both organophosphate insecticides tested (pirimiphos-methyl and chlorpyriphos-methyl). No mortality (0%) was observed in the negative control tubes (silicon oil). Mortality rate of the susceptible *An. gambiae s.s*. Kisumu mosquitoes for all the insecticides tested was 100%.

**Fig 1.**
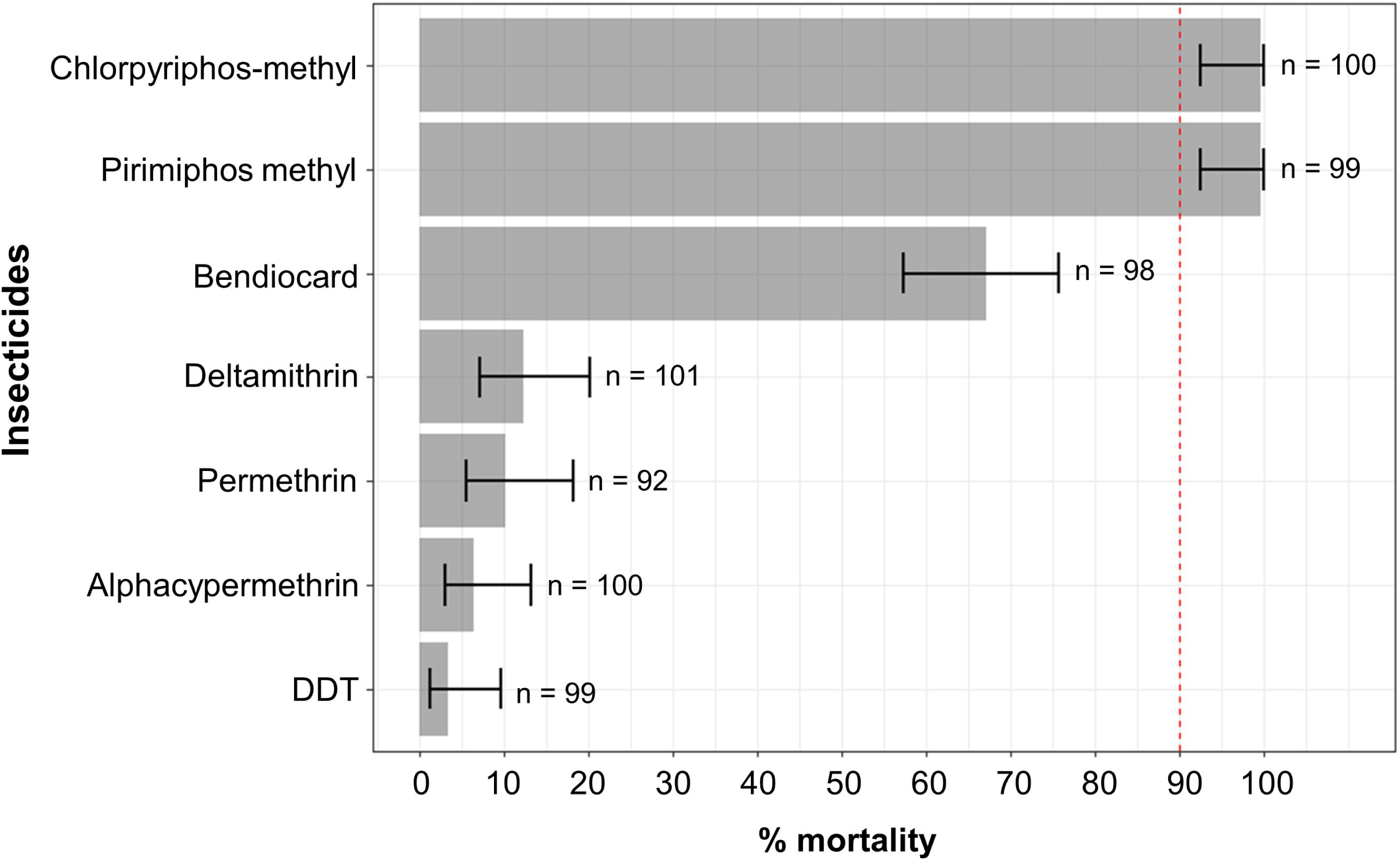
Susceptibility of wild *An. gambiae s.l.* from southwest Burkina Faso to seven insecticides used for malaria control. Bars indicate estimated marginal means (EMM) of mortality as predicted by a generalized linear model. Error bars represent the 95% confidence intervals of the EMMs. If mortality falls under 90% (red dashed line), the mosquito population is considered resistant to the tested insecticide.

### Chemical analysis

On cement walls, chemical analysis indicates that the mean concentrations of pirimiphos-methyl on classical and plasticized filter papers were 1428 mg/m^2^ (CI95% [719; 2136]) and 1421 mg/m^2^ (CI95% [713; 2130]), respectively (Fig. 2A). We did not find a difference in pirimiphos-methyl concentration between the classical and plasticized papers applied on cement walls (mean difference (MD) = 6.13 mg/m^2^ (CI95% [-309; 322]), Tukey’s test p-value = 0.96).

**Fig. 2.**
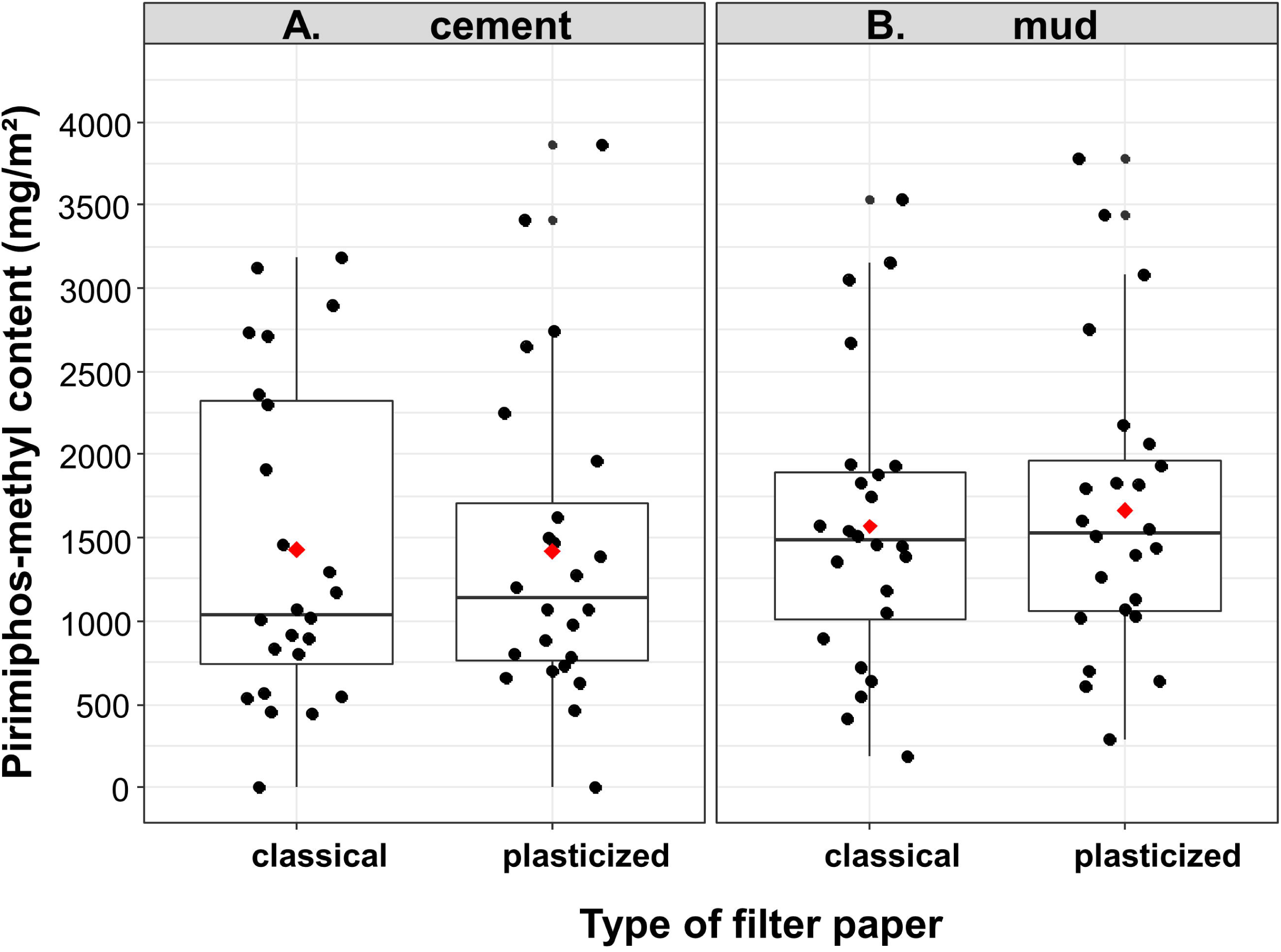
Applied doses of pirimiphos-methyl on cement (A) and mud (B) walls. Red diamonds show the mean concentrations of pirimiphos-methyl on filter papers. Boxes show 1^st^ and 3^rd^ quartiles as well as the median concentration. The whiskers extends to the largest and lowest values that are no further than 1.5 * IQR (where IQR is the inter-quartile range, or distance between the first and third quartiles). Black dots represent concentration measured for all filter papers.

On mud walls, chemical analysis indicates that the mean concentrations of pirimiphos-methyl on classical and plasticized papers from sprayed houses were 1569 mg/m^2^ (CI95% [861; 2278]) and 1665 mg/m^2^ (CI95% [957; 2373]), respectively (Fig. 2B). We were not able to find a difference in pirimiphos-methyl concentration between classical and plasticized papers applied on mud walls (MD = -95.76, CI95% [-411; 220], Tukey’s test p-value=0.54). Moreover, pirimiphos-methyl concentration on papers placed on cement did not differ from that on mud walls (MD = -193, CI95% [-1039; 653], Tukey’s test p-value =0.65).

### Insecticide residual efficacy

Predictions from the mortality model of *An. gambiae s.l* wild strain showed that pirimiphos-methyl efficacy remained higher than 80% until the last test (i.e. 210 days after spraying) on both cement and mud walls (Fig. 3A). We were not able to find a difference in residual efficacy of pirimiphos-methyl between cement and mud walls with *An. gambiae s.l* wild strain (OR = 0.792, CI95% [0.55; 1.12], p-value =0.19).

**Fig. 3.**
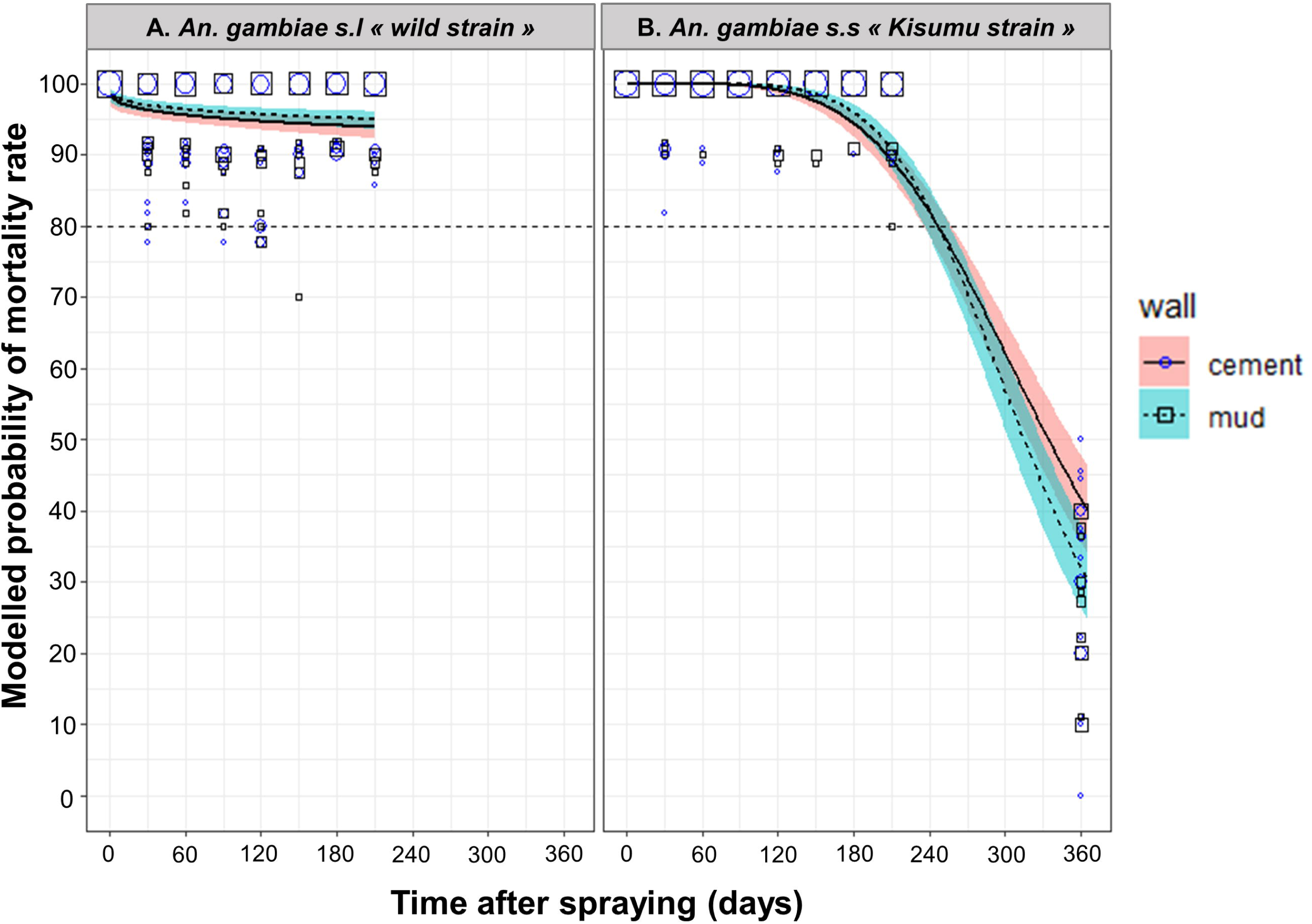
Efficacy (mortality) over time of indoor residual spraying of pirimiphos-methyl against wild *An. gambiae s.l.* (A) and susceptible *An. gambiae s.s* (B). Mortality rates were predicted from a binomial-response mixed effect model. Pirimiphos-methyl at 4 ml/m^2^ targeted dose applied on mud (dashed lines) or cement (solid lines) walls are compared. Grey areas are 95% confidence interval of predicted means. Mortality values measured on the field are shown as blue circles (cement) and black squares (mud) of size proportional to the number of values (max = 20).

With the susceptible *An. gambiae s.s. kisumu* strain, for which a supplementary test was done on day 360 post-spraying, predictions from the mortality model show that pirimiphos-methyl treatment was effective (mortality >80%) until the 247^th^ day post-spraying, both on cement and mud walls (Fig. 3B). The residual efficacy of pirimiphos-methyl was lower on mud than on cement walls (OR=0.257, CI95% [0.07; 0.86], p-value =0.02).

## Discussion

Insecticide resistance management has remained a major challenge for malaria control and elimination for years (WHO 2012, 2018b). This is, in large part, due to the fact that malaria vectors are developing resistance to most of the insecticides currently used in public health (WHO 2018b). Insecticide susceptibility assays showed high resistance of *An. gambiae s.l.* from Diébougou to all pyrethroids tested (deltamethrin, permethrin and alphacypermethrin). Our results are consistent with that of a recent investigation conducted in an area of southwest Burkina Faso in 2016 (Namountougou et al. 2019). But, compared with data collected pre-2010, this study suggests that the prevalence of pyrethroids resistance have increased considerably over time (Dabiré et al. 2009, 2012, Badolo et al. 2012). In addition, DDT and Bendiocarb induced respectively 4% and 67% mortality rates indicating a multi-resistance of the wild *An. gambiae s.l* populations to pyrethroids, organochlorines and carbamates.

Many mechanisms might be involved in this multiple resistance. Indeed, both L1014F and L1014S *kdr* mutations that confer cross-resistance to organochlorine and pyrethroids were recorded in a neighboring population from Diébougou (Namountougou et al. 2019). The same authors also describe the presence at low frequency of the *ace-1* mutation that confer cross-resistance to both carbamates and organophosphorous and evidenced the presence of metabolic resistance mechanisms (esterase and GST) that may confer resistance to all tested families of insecticides (WHO 2018b). The large-scale use of LLINs across the country (Protopopoff et al. 2008) might have contributed to the selection of these resistance mechanisms, particularly those involved in pyrethroid resistance as well as the intense use of insecticides in agriculture (Diabate et al. 2002, Yadouleton et al. 2011, Hien et al. 2017). Nevertheless, the wild population of *An. gambiae s.l.* from our study area was found to be fully susceptible to organophosphorus compounds (chlorpyrifos-methyl and pirimiphos-methyl). These data were strengthened by the results of the WHO cone bioassay done in houses of two villages sprayed with pirimiphos-methyl CS. Indeed, the duration of residual efficacy (mortality >80%) of pirimiphos-methyl IRS on mud and cement walls was longer than 7 months against wild strains of *An. gambiae*. Unfortunately, we were not able to determine the precise effective duration because further testing was not performed beyond 7 months.

The mortality model predicted that the residual efficacy of pirimiphos-methyl IRS lasted for 247 days (8-9 months) against the susceptible *An. gambiae* Kisumu strain. In Benin, pirimiphos-methyl sprayed in experimental huts has shown 9 and 6 months of effective residual efficacy on cement and mud substrates, respectively, against susceptible *An. Gambiae* Kisumu (Rowland et al. 2013). However, these durations dropped to 5 months on both substrates in houses of northern Benin with susceptible *An. gambiae* Kisumu (Salako et al. 2019). In Ivory Coast, 5 and 7 months residual efficacy were observed on mud and cement walls, respectively, against *An. gambiae* Kisumu (Tchicaya et al. 2014). In Ethiopia, Yewhalaw and colleagues observed a 6 months residual efficacy against a susceptible strain of *An. arabiensis* on mud substrates (Yewhalaw et al. 2017). In Tanzania, pirimiphos-methyl displayed 3 to 6 months of residual efficacy depending on the substrate. In a multi-country study (Dengela et al. 2018), pirimiphos-methyl CS duration of residual efficacy ranged from 2 to 9 months. Many factors such as quality of spraying (Fuseini et al. 2020), vector resistance (Ranson and Lissenden 2016, WHO 2018b), season/climate (Parham and Hughes 2015) and wall modifications post-application (Opiyo and Paaijmans 2020) can explain the observed differences in residual efficacy between sites and studies.

In this study, we did not found any difference in the residual efficacy between cement and mud surfaces. However, according to the results of the above-mentioned studies, insecticides often performed poorly on mud surfaces probably because these surfaces are porous and hence absorb a quantity of the applied insecticide. Our results suggest that mud walls in the rural area of Diébougou might be less porous. The absence of difference in the residual efficacy between cement and mud surfaces could also be explained by the quantity of insecticides sprayed on the walls which exceed the recommended dose. Indeed, chemical analysis of classical filter papers showed that applied to target dose ratios were 1.43 [0.81; 2.05] on cement surfaces and 1.57 [0.95; 2.19] on mud surfaces. This would indicate that the residual efficacy of pirimiphos-methyl CS would be reduced if the applied dose was within the recommended ± 25% limit of the target dose. In this study, we have tested two filter papers: classical papers and plasticized papers. Plasticized papers were tested because Moiroux *et al.* (Moiroux et al. 2018) found that the concentration of alpha-cypermethrin on filter paper placed on mud walls was lower than on filter papers placed on cement wall. They hypothesized that a quantity of the insecticide applied on mud surfaces may migrate from the filter papers to the wall, diminishing the insecticide content on classical papers placed on mud surfaces. Consequently, they suggest the use of plasticized papers to address the issue (Moiroux et al. 2018). Here, we did not find any differences in concentration between papers placed on mud or cement surface. Further replications of this experiment are nevertheless recommended as we did not use the same insecticide as Moiroux *et al*.

To date, only microencapsulated formulation of pirimiphos-methyl and new formulations of clothianidin (neonicotinoids) (SumiShield^®^ 50WG and Fludora Fusion^®^ WP-SB) have the potential to be effective for more than 6 months (WHO 2015, 2019b). That was confirmed in this study for pirimiphos-methyl. For clothianidin, no field trial was carried out to evaluate its residual efficacy as IRS in southwest Burkina Faso but local vector populations were shown to be susceptible (Oxborough et al. 2019). Therefore, pirimiphos-methyl and clothianidin used in rotation or mosaic might constitute an effective insecticide resistance management strategy in southwest Burkina Faso.

## Conclusions

The *Anopheles gambiae s.l.* population from the rural area of Diébougou, in the southwest of Burkina Faso was resistant to DDT, all pyrethroids tested and bendiocarb. In contrast the same population was susceptible to both OPs tested (pirimiphos-methyl and chlorpyrimiphos-methyl). This result was further supported by the residual efficacy of pirimiphos-methyl IRS which has lasted more than 7 months on cement and mud walls against both susceptible *An. gambiae s.s* kisumu strain and wild *An. gambiae s.l.* populations. In the context of insecticide resistance management, pirimiphos-methyl CS could be integrated in IRS and used in rotation with others families of insecticides that have been shown to be effective against the local malaria vector populations.

## Acknowledgements

We are grateful to the Burkina Faso Ministry of Health and local authorities for their participation in the study. We acknowledge the logistic support (spray pumps) provided to the project by National Malaria Control Programs. We thank the Diébougou district medical team, particularly Dr. Dembélé Henri and Mr. Kaboré Adama for field support during insecticide application. We thank the technical staff of IRSS-DRO especially to Ouari Ali, Meda G. Benson and Millogo S. Abel for their assistance. We also thank to Ouattara Adama and Dahounto Amal for their support. We extend our sincere appreciation to the populations from all selected villages, spray operators and all volunteers for the support and kind cooperation during fieldwork implementation.

This work was part of the REACT project, funded by the French Initiative 5% – Expertise France (No. 15SANIN213). NM, RKD and DDS conceived and designed the study. DDS, FDH, ASH, DPAK and MK collected the data. DDS analyzed the data. DDS and BZ drafted the manuscript. CP, GAO, DPAK, MK, AAK, NM and RKD reviewed the manuscript; all authors read and approved the final manuscript. The authors declare that they have no competing interests.

## References Cited

Badolo, A., A. Traore, C. M. Jones, A. Sanou, L. Flood, W. M. Guelbeogo, H. Ranson, and N. F. Sagnon. 2012. Three years of insecticide resistance monitoring in *Anopheles gambiae* in Burkina Faso□: resistance on the rise□? Malar. J. 11: 1–11.

Bhatt, S., D. J. Weiss, E. Cameron, D. Bisanzio, B. Mappin, U. Dalrymple, K. E. Battle, C. L. Moyes, A. Henry, P. A. Eckhoff, E. A. Wenger, O. Briët, M. A. Penny, T. A. Smith, A. Bennett, J. Yukich, T. P. Eisele, J. T. Griffin, C. A. Fergus, M. Lynch, F. Lindgren, J. M. Cohen, C. L. J. Murray, D. L. Smith, S. I. Hay, R. E. Cibulskis, and P. W. Gething. 2015. The effect of malaria control on *Plasmodium falciparum* in Africa between 2000 and 2015. Nature. 526: 207–211.

Dabiré, K., A. Diabaté, M. Namountougou, L. Djogbenou, C. Wondji, F. Chandre, F. Simard, J. Ouédraogo, T. Martin, M. Weill, and T. Baldet. 2012. Trends in insecticide resistance in natural populations on malaria vectors in Burkina Faso, West Africa: 10 Years’ surveys. Insectic. - Pest Eng. 479–502.

Dabiré, K. R., A. Diabaté, M. Namountougou, K. H. Toé, A. Ouari, P. Kengne, C. Bass, and T. Baldet. 2009. Distribution of pyrethroid and DDT resistance and the L1014F kdr mutation in *Anopheles gambiae* s.l. from Burkina Faso (West Africa). Trans. R. Soc. Trop. Med. Hyg. 103: 1113–1120.

Dengela, D., A. Seyoum, B. Lucas, B. Johns, K. George, A. Belemvire, A. Caranci, L. C. Norris, and C. M. Fornadel. 2018. Multi-country assessment of residual bio-efficacy of insecticides used for indoor residual spraying in malaria control on different surface types: Results from program monitoring in 17 PMI/USAID-supported IRS countries. Parasit. Vectors. 11: 1–14.

Diabate, A., T. Baldet, F. Chandre, M. Akogbeto, T. R. Guiguemde, F. Darriet, C. Brengues, P. Guillet, J. Hemingway, G. J. Small, and J. M. Hougard. 2002. The role of agricultural use of insecticides in resistance to pyrethroids in *Anopheles gambiae* s.l. in Burkina Faso. Am. J. Trop. Med. Hyg. 67: 617–622.

Djènontin, A., O. Aïmihouè, M. Sèzonlin, G. B. Damien, R. Ossè, B. Soukou, G. Padonou, F. Chandre, and M. Akogbéto. 2013. The residual life of bendiocarb on different substrates under laboratory and field conditions in Benin, Western Africa. BMC Res. Notes. 6: 1–6.

Firth, D. 1993. Amendments and corrections: bias reduction of maximum likelihood estimates. Biometrika. 80: 27–38.

Fuseini, G., H. M. Ismail, M. E. Von Fricken, T. A. Weppelmann, J. Smith, R. A. Ellis Logan, F. Oladepo, K. J. Walker, W. P. Phiri, M. J. I. Paine, and G. A. Garciá. 2020. Improving the performance of spray operators through monitoring and evaluation of insecticide concentrations of pirimiphos-methyl during indoor residual spraying for malaria control on Bioko Island. Malar. J. 19: 1–8.

Hien, S. A., D. D. Soma, O. Hema, B. Bayili, M. Namountougou, O. Gnankiné, T. Baldet, A. Diabaté, and K. R. Dabiré. 2017. Evidence that agricultural use of pesticides selects pyrethroid resistance within *Anopheles gambiae* s.l. populations from cotton growing areas in Burkina Faso, West Africa. PLoS One. 3: 1–15.

Institut National de la Statistique et de la Démographie. 2015. Tableau de bord économique et social 2014 de la région du Sud-Ouest. : 1–64.

Institut National de la Statistique et de la Démographie. 2017. Enquête nationale sur le secteur de l’orpaillage (ENSO). : 1–10.

Kosmidis, I. 2019. Bias reduction in binomial-response generalized linear models. pp. 1–24. https://github.com/ikosmidis/brglm BugReports.

Moiroux, N., A. Djènontin, B. Zogo, A. Bouraima, I. Sidick, and O. Pigeon. 2018. Small-scale field testing of alpha-cypermethrin water-dispersible granules in comparison with the recommended wettable powder formulation for indoor residual spraying against malaria vectors in Benin. Parasit. Vectors. 11: 1–8.

Namountougou, M., D. D. Soma, M. Kientega, M. Balboné, D. P. A. Kaboré, S. F. Drabo, A. Y. Coulibaly, F. Fournet, T. Baldet, A. Diabaté, R. K. Dabiré, and O. Gnankiné. 2019. Insecticide resistance mechanisms in *Anopheles gambiae* complex populations from Burkina Faso, West Africa. Acta Trop. 197: 1–9.

Opiyo, M. A., and K. P. Paaijmans. 2020. “We spray and walk away”: Wall modifications decrease the impact of indoor residual spray campaigns through reductions in post-spray coverage. Malar. J. 19: 1–6.

Oxborough, R. M., A. Seyoum, Y. Yihdego, R. Dabire, V. Gnanguenon, F. Wat’senga, F. R. Agossa, G. Yohannes, S. Coleman, L. M. Samdi, A. Diop, O. Faye, S. Magesa, A. Manjurano, M. Okia, E. Alyko, H. Masendu, I. Baber, A. Sovi, J.-D. Rakotoson, K. Varela, B. Abong’o, B. Lucas, C. Fornadel, and D. Dengela. 2019. Susceptibility testing of Anopheles malaria vectors with the neonicotinoid insecticide clothianidin; results from 16 African countries, in preparation for indoor residual spraying with new insecticide formulations. Malar. J. 18: 1–11.

Parham, P. E., and D. A. Hughes. 2015. Climate influences on the cost-effectiveness of vector-based interventions against malaria in elimination scenarios. Philos. Trans. R. Soc. B Biol. Sci. 370: 1–15.

Poshadri, A, and A. Kuna. 2010. Microencapsulation technology: A review. J. Res. ANGRAU. 38: 86–102.

Protopopoff, N., W. Van Bortel, T. Marcotty, M. Van Herp, P. Maes, D. Baza, U. D’Alessandro, and M. Coosemans. 2008. Spatial targeted vector control is able to reduce malaria prevalence in the highlands of Burundi. Am. J. Trop. Med. Hyg. 79: 12–18.

Ranson, H., and N. Lissenden. 2016. Insecticide resistance in African *Anopheles* mosquitoes: A worsening situation that needs urgent action to maintain malaria control. Trends Parasitol. 32: 187–196.

Rowland, M., P. Boko, A. Odjo, A. Asidi, M. Akogbeto, and R. N’Guessan. 2013. A new long-lasting indoor residual formulation of the organophosphate insecticide pirimiphos methyl for prolonged control of pyrethroid-resistant mosquitoes: An experimental hut trial in Benin. PLoS One. 8: 1–10.

Russell, L., S. Henrik, J. Love, P. Buerkner, and M. Herve. 2019. estimated marginal means, aka least-squares means: package ‘emmeans.’ https://github.com/rvlenth/emmeans.

Salako, A. S., F. Dagnon, A. Sovi, G. G. Padonou, R. Aïkpon, I. Ahogni, T. Syme, R. Govoétchan, H. Sagbohan, A. A. Sominahouin, B. Akinro, L. Iyikirenga, F. Agossa, and M. C. Akogbeto. 2019. Efficacy of Actellic 300 CS-based indoor residual spraying on key entomological indicators of malaria transmission in Alibori and Donga, two regions of northern Benin. Parasit. Vectors. 12: 1–14.

Sibanda, M. M., W. W. Focke, F. J. W. J. Labuschagne, L. Moyo, N. S. Nhlapo, A. Maity, H. Muiambo, P. M. Jr, N. A. S. Crowther, M. Coetzee, and G. W. A. Brindley. 2011. Degradation of insecticides used for indoor spraying in malaria control and possible solutions. Malar. J. 10: 1–12.

Soma, D. D., B. M. Zogo, A. Somé, B. N. Tchiekoi, D. F. de S. Hien, H. S. Pooda, S. Coulibaly, J. E. Gnambani, A. Ouari, K. Mouline, A. Dahounto, G. A. Ouédraogo, F. Fournet, A. A. Koffi, C. Pennetier, N. Moiroux, and R. K. Dabiré. 2019. *Anopheles* bionomics, insecticide resistance and malaria transmission in southwest Burkina Faso□: a pre-intervention study. BioRxiv. 17: 1–39.

Tchicaya, E. S., C. Nsanzabana, T. A. Smith, J. Donzé, M. de Hipsl, Y. Tano, P. Müller, O. J. Briët, J. Utzinger, and B. G. Koudou. 2014. Micro-encapsulated pirimiphos-methyl shows high insecticidal efficacy and long residual activity against pyrethroid-resistant malaria vectors in central Côte d’Ivoire. Malar. J. 13: 1–13.

The R Development Core Team. 2008. R□: A Language and Environment for Statistical Computing. : 1–2547.

World Health Organization (WHO). 1969. Re-examination of the Global Strategy of Malaria Eradication Official records of the World Health Organization: Twenty-second World Health Assembly. World Heal. Organ. Geneva, Switzerland. : 106–126.

World Health Organization (WHO). 2006. Pesticides and their application for the control of vectors and pests of public health importance. World Heal. Organ. Geneva, Switzerland. 1: 1–104.

World Health Organization (WHO). 2007. Manuel for indoor residual spraying□: Application of residual sprays for vector control Third edition. World Heal. Organ. Geneva, Switzerland. : 1–53.

World Health Organization (WHO). 2011. The use of DDT in malaria vector control. Global malaria programme. World Heal. Organ. Geneva, Switzerland. : 1–9.

World Health Organization (WHO). 2012. Global plan for insecticide resistance management in malaria vectors. World Heal. Organ. Geneva, Switzerland. : 1–130.

World Health Organization (WHO). 2015. World malaria report 2015, World Heal. Organ. Geneva, Switzerland. : 1–181.

World Health Organization. 2016. Test procedures for insecticide resistance monitoring in malaria vector mosquitoes Second edition. World Heal. Organ. Geneva, Switzerland. : 1–48.

World Health Organization (WHO). 2018a. World malaria report 2018. World Heal. Organ. Geneva, Switzerland. : 1-238.

World Health Organization (WHO). 2018b. Global report on insecticide resistance in malaria vectors: 2010–2016. World Heal. Organ. Geneva, Switzerland. : 1–72.

World Health Organization (WHO). 2019a. World malaria report 2019. World Heal. Organ. Geneva, Switzerland. : 1–185.

World Health Organization (WHO). 2019b. List of WHO Prequalification vector control products. World Heal. Organ. Geneva, Switzerland. :1–5.

Yadouleton, A., T. Martin, G. Padonou, F. Chandre, A. Asidi, L. Djogbenou, R. Dabiré, R. Aïkpon, M. Boko, I. Glitho, and M. Akogbeto. 2011. Cotton pest management practices and the selection of pyrethroid resistance in *Anopheles gambiae* population in Northern Benin. Parasit. Vectors. 4: 1–11.

Yewhalaw, D., M. Balkew, J. Shililu, S. Suleman, A. Getachew, G. Ashenbo, S. Chibsa, G. Dissanayake, K. George, D. Dengela, Y. Ye-Ebiyo, and S. R. Irish. 2017. Determination of the residual efficacy of carbamate and organophosphate insecticides used for indoor residual spraying for malaria control in Ethiopia. Malar. J. 16: 1–9.

Zhao, Z.-Q., Z.-G. Yu, V. Anh, J.-Y. Wu, and G.-S. Han. 2016. Protein folding kinetic order prediction from amino acid sequence based on horizontal visibility network. Curr. Bioinform. 11: 173–185.

